# Age-Associated Remodeling of neuroglial Connexin43 in the Mouse and Human Brain

**DOI:** 10.64898/2026.05.18.725990

**Authors:** Qi Wang, Chenmeng Wang, Hui Li, Mengjiao Sun, Wei-na Jin, Alexei Verkhratsky, Chenju Yi

## Abstract

Age-dependent changes in the major neuroglial junctional channel Connexin43 (Cx43) are poorly defined. We integrated public multi-organ and single-cell datasets with high-resolution morphological, biochemical, and functional analyses of human and mouse brains. We reveal a non-linear aging trajectory for Cx43, marked by an adaptational midlife elevation followed by a late-life decline. These molecular changes proceed in the absence of significant cell loss but are associated with extensive glial remodeling. Notably, while astrocytic gap junction coupling in the hippocampus remains largely preserved, aging induces a selective increase in hemichannel mediated dye uptake in both microglia and hippocampal astrocytes. In human cortical and hippocampal tissues, we confirm that aging drives a significant reduction in astrocytic Cx43 expression, alongside characteristic morphological simplification of cell structure and domain contraction. Together, our findings redefine the aging brain as a state of active, multi-level glial remodeling rather than a simple decline in connexin-mediated communication.

## 1. INTRODUCTION

Aging population is the most pressing worldwide public health challenges [1]. Aging is the major risk factor for cognitive decline and for neurodegenerative disorders such as Alzheimer’s disease and Parkinson’s disease [2, 3]. Although neuronal vulnerability has long been central to studies of brain aging and age-dependent disorders, glial dysfunction is increasingly recognized as a fundamental contributor [4-8]. Astrocytes are key homeostatic neuroglia supporting neurotransmitter recycling, metabolic homeostasis, neurovascular coupling, and blood–brain barrier (BBB) integrity as well as to contributing to multiple aspects of information processing in the CNS [9-13]. Aging reshapes astrocyte transcriptional states and is associated with astroglial atrophy across brain regions in humans and rodents [14-19]. Even so, the extent, regional specificity, and functional consequences of astrocyte aging remain incompletely resolved.

A central feature of astrocyte biology is their extensive homocellular (astrocyte-astrocyte) and heterocellular (astrocyte-oligodenrocyte/ependymoglia) communication networks [20]. Through gap junctions and/or hemichannels, astrocytes help maintain ionic, metabolic, and signaling balance that is essential for neuronal survival and circuit stability [21, 22]. These channels are formed by connexins, with Connexin43 (Cx43) being the predominant astrocytic connexin in the brain [23]. Cx43 is therefore well positioned to influence how astrocytic coordination of tissue-level homeostasis under both physiological and pathological conditions [24]. Indeed, Cx43 dysregulation has been implicated in neurodegenerative, neuropsychiatric, and inflammatory disorders [25-29]. However, its specific role and temporal dynamics during normal brain aging remain poorly defined.

A coherent view on Cx43 changes in the aging brain remains to be elucidated. Reduced astrocytic gap junction coupling was detected in older animals, even when total protein abundance appears relatively stable, suggesting structural remodeling of astrocytic networks rather than a simple loss of expression [30]. By contrast, increased Cx43 expression and enhanced hemichannel opening is frequently reported in reactive astrocytes in neuroinflammation, neuronal injury, and BBB dysfunction [31, 32]. These findings are informative, but they do not resolve what happens during normal aging itself. Moreover, many prior studies have relied on bulk tissue measurements or disease models and therefore lack the spatial and cell-state resolution needed to capture heterogeneous astrocyte responses across the aging brain. Taken together, these data leave a central question unresolved: does brain aging lead to a uniform decline in astrocytic gap junction coupling, or does it drive a region- and state-specific reorganization of Cx43-dependent network architecture that shapes neuronal vulnerability or resilience?

Here, we address this question by combining single-cell and histological approaches to examine Cx43 in the aging brain. We aim to define how age-related changes in Cx43 expression, localization, and channel function alter intercellular communication and influence neural vulnerability. Resolving this issue is important not only for understanding normal brain aging, but also for defining glial remodeling on which age-dependent neurodegenerative diseases develop. Distinguishing adaptive from maladaptive Cx43 remodeling has become an urgent step toward understanding, and eventually targeting, early mechanisms of brain vulnerability.

## 2. RESULTS

### 2.1 Cx43 is enriched in astrocytes throughout brain aging

To characterize the tissue and cellular landscape of Cx43 during aging, we re-analyzed published mouse multi-organ aging datasets [33, 34]. Our analysis of a multi-organ proteomic dataset revealed that Cx43 is consistently more abundant in the brain than in peripheral tissues, including the heart, intestine, lung, skin, and spleen (Fig. 1A). Furthermore, brain *Gja1* expression is maintained across the lifespan, showing no evidence of any decline with age (Fig. 1B). To identify the cellular source of brain *Gja1*, we interrogated a published mouse brain single-cell atlas [35]. *Gja1* expression was strongly enriched in astrocytes, with negligible expression in other glial, vascular, or neuronal populations (Fig. 1C). This expression pattern was recapitulated in a human non-neuronal brain single-cell atlas, where *GJA1* was predominantly detected in astrocytes while remaining detectable in other non-neuronal cell types (Fig. 1D) [36]. Together, these findings demonstrate that brain Cx43 expression is broadly maintained during aging and remains preferentially associated with astrocytes. This transcript-level stability provided a reference point for examining whether Cx43 protein abundance and distribution are similarly preserved during brain aging.

**Figure 1.**
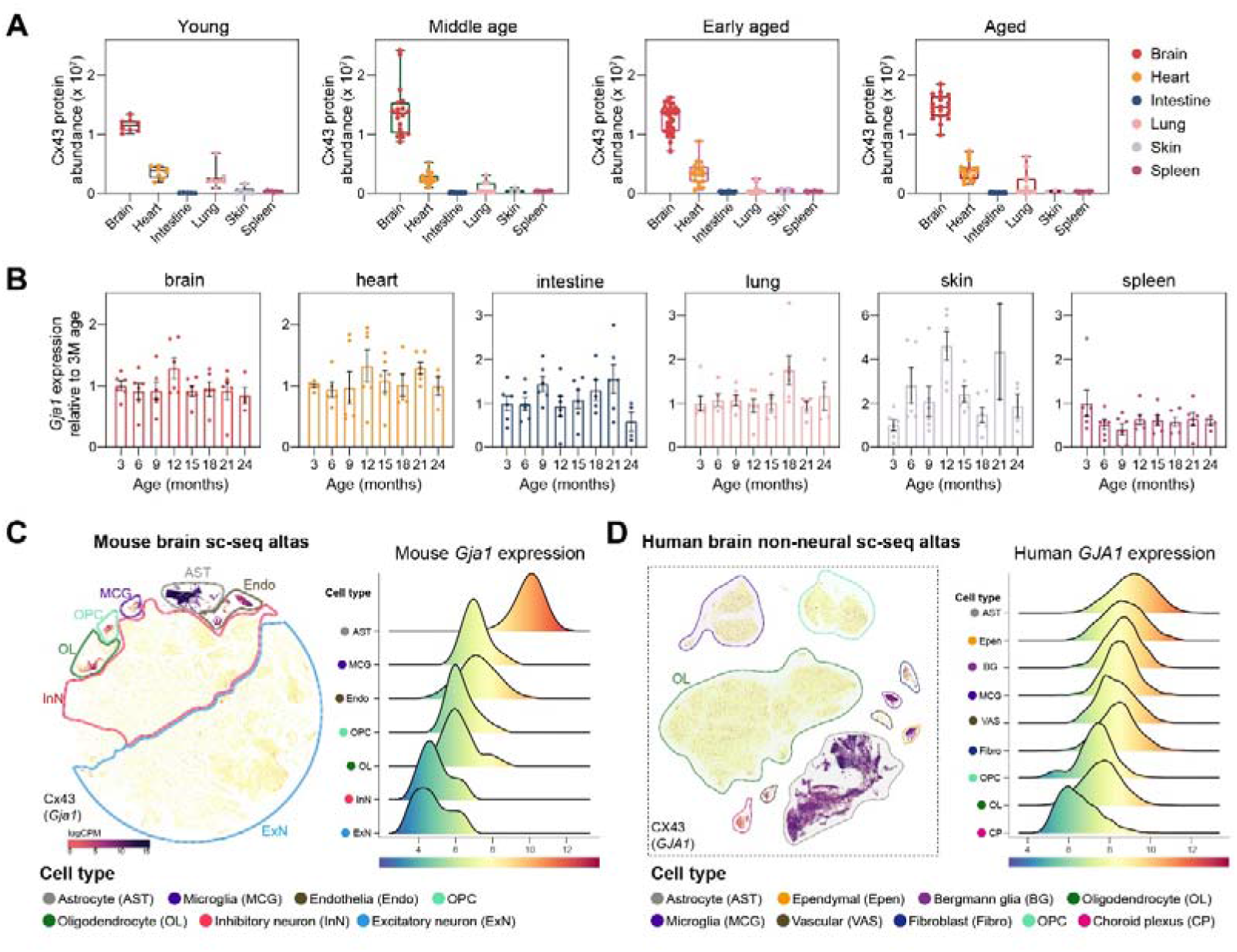
Cx43 is enriched in the brain and preferentially associated with astrocyte-centered brain cellular networks. **(A)** Cx43 protein abundance across mouse organs at different life stages (young, middle age, early aged, and aged). **(B)** Relative *Gja1* expression across age in mouse brain, heart, intestine, lung, skin, and spleen. **(C)** Mouse brain single-cell atlas showing cell-type distribution of *Gja1* expression. Left, UMAP of major brain cell populations. Right, ridge plots of *Gja1* expression across annotated cell types. **(D)** Human brain non-neural single-cell atlas showing cell-type distribution of *GJA1* expression. Left, UMAP of major non-neuronal cell populations. Right, ridge plots of *GJA1* expression across annotated cell types.

### 2.2 Cx43 expression is dynamically remodeled in the aging mouse brain independent of cell loss

We next asked whether the stable brain *Gja1* mRNA pattern was mirrored at the protein level. To address this, we examined the age-related dynamics of Cx43 in the mouse brain across four time points (4, 12, 18, and 24 months). Whole-brain immunofluorescence analysis demonstrated robust Cx43 expression in the cortex and hippocampus. Regional heatmapping identified a non-linear temporal pattern, with an elevation of Cx43 levels at 12 months followed by a subsequent decline at 18 and 24 months. Thus, despite broadly maintained *Gja1* expression at the tissue level, brain Cx43 protein undergoes region-dependent remodeling throughout the aging process (Fig. 2A).

**Figure 2.**
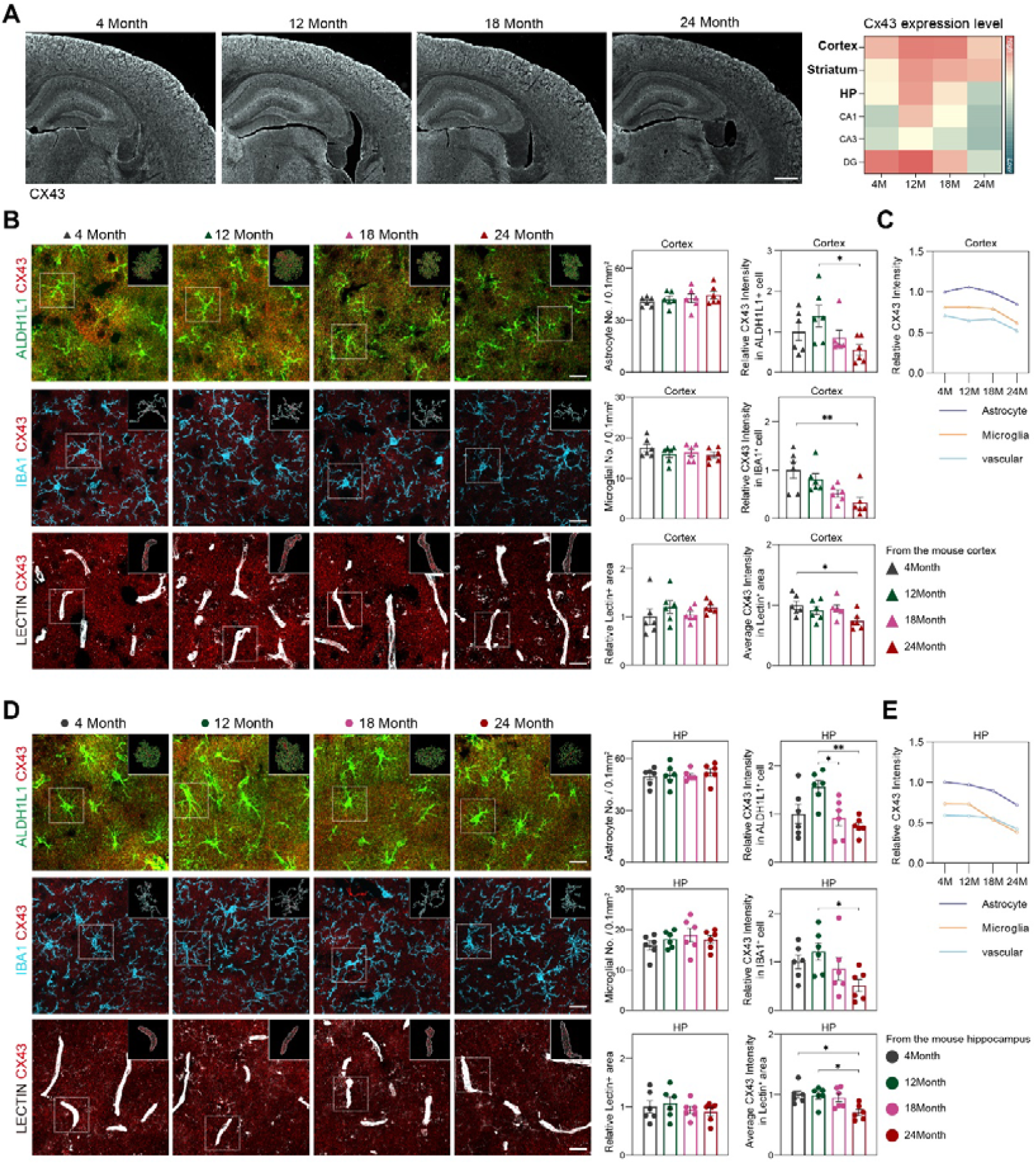
Aging remodels Cx43 expression and cellular distribution in the mouse cortex and hippocampus. **(A)** Representative brain wide Cx43 immunofluorescence images from mice aged 4, 12, 18, and 24 months. Heatmap showing relative Cx43 expression across brain regions. **(B, C)** Representative cortical images co-stained for Cx43 with ALDH1L1, IBA1, or Lectin across age groups, with corresponding quantification. **(D, E)** Representative hippocampal images co-stained for Cx43 with ALDH1L1, IBA1, or Lectin across age groups, with corresponding quantification. Data are shown as mean ± SEM. Each dot represents one biological replicate. Scale bar represents 500 μm in (A) and 20 μm in (B, D).

To quantify Cx43 expression in different brain cells we utilized specific markers— ALDH1L1 for astrocytes, IBA1 for microglia, and lectin for microvessels. Quantification confirmed that the densities of astrocytes, microglia, and vascular area remained constant in the cortex across all age groups, indicating that the observed Cx43 fluctuations are not due to cell loss (Fig. 2B). Cx43 immunoreactivity in different cells changed significantly: astrocytic Cx43 expression peaked at 12 months, whereas Cx43 signal associated with microglia and vasculature declined during later stages of aging (Fig. 2B). Comparisons across all time points conformed that astrocytes represent the primary reservoir of Cx43 in the brain (Fig. 2C).

A similar pattern was observed in the hippocampus (Fig. 2D). While glial cell abundance and vascular coverage remained stable, Cx43 intensity showed clear age-dependent reorganization. Hippocampal astrocytes exhibited the characteristic ‘mid-life peak’ followed by a late-life decline, whereas Cx43 reduction was most pronounced in microglial and vascular compartments of aged mice (Fig. 2D). These findings demonstrate that the aging brain is characterized by a time-controlled, compartment-specific regulation of Cx43, while highlighting astrocytes as the central hub of this remodeling process.

### 2.3 Aging drives region-specific remodeling of astrocyte and microglial architecture

The age-dependent changes in cell-associated Cx43 expression raised the question of whether glial structure is similarly compromised during aging. To address this, we performed 3D reconstruction of individual astrocytes and microglia in the cortex and hippocampus, utilizing Sholl analysis, territorial domain analysis, and soma-to-territory ratio quantification.

Cortical astrocytes showed preserved morphology between 4 and 12 months, followed by structural simplification at 18 and 24 months. This decline was reflected by reduced branching complexity, smaller territorial domains, and an increased soma-to-territory ratio (Fig. 3A). In addition, volume fraction analysis revealed a reduction in the astrocytic process-rich domain at 18 and 24 months, indicating that cortical astrocyte remodeling involved not only major branch retraction but also loss of fine peripheral processes (Fig. 3B). Hippocampal astrocytes, however, followed a more distinct non-linear trajectory: they exhibited increased complexity and territorial expansion at 12 months, followed by marked contraction during later stages of aging (Fig. 3C). Consistently, volume fraction analysis showed a decline in astrocytic process occupancy in the aged hippocampus, further supporting an age-associated reduction of the fine astrocytic network (Fig. 3D).

**Figure 3.**
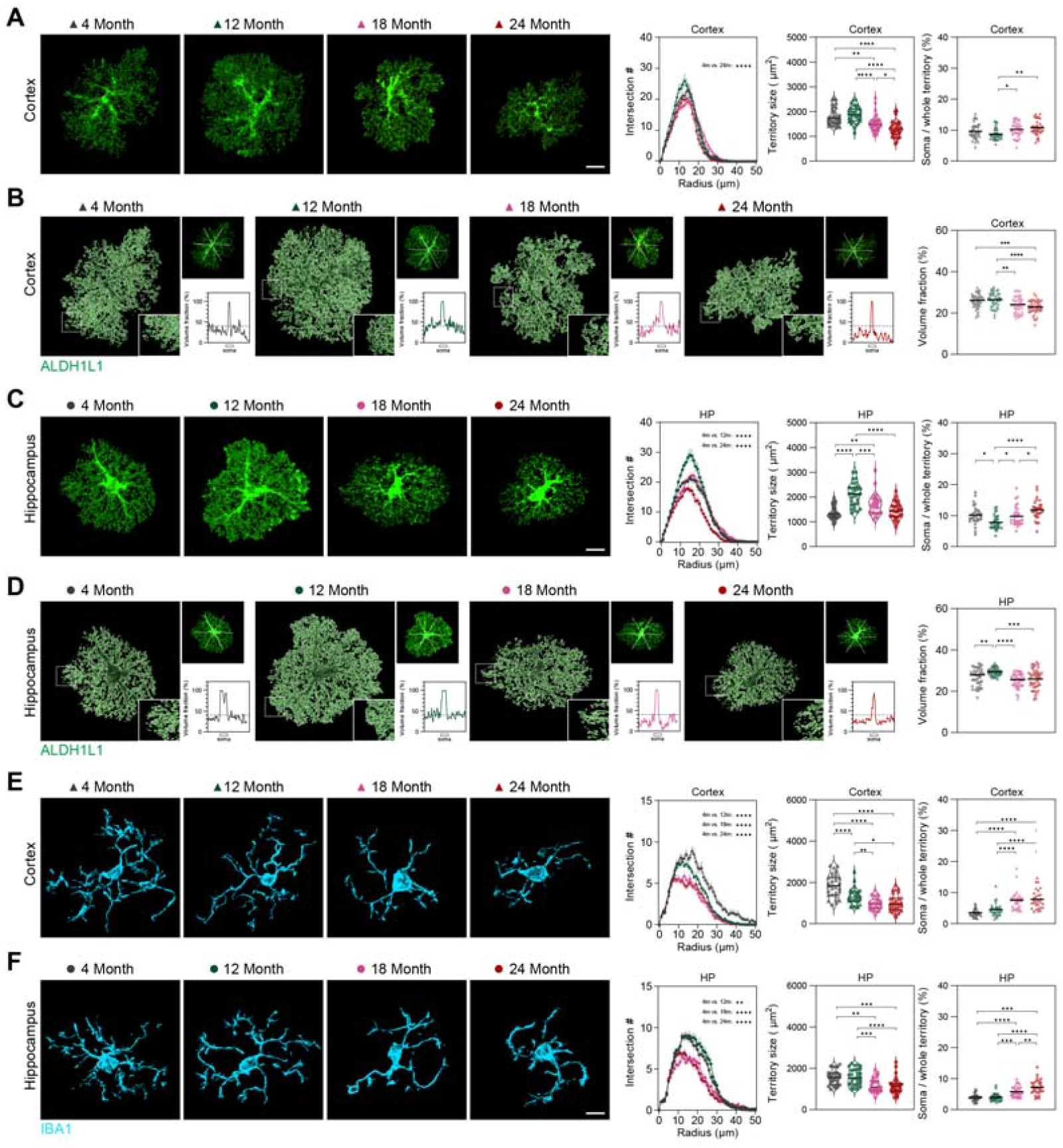
Aging remodels astrocyte and microglial morphology in the mouse cortex and hippocampus. **(A, C)** Representative reconstructions of ALDH1L1+ astrocytes in the cortex (A) and hippocampus (C) from mice aged 4, 12, 18, and 24 months, with corresponding quantification of Sholl intersections, territory size, and soma-to-whole territory ratio. **(B, D)** Representative reconstructions illustrating astrocytic volume fraction in the cortex (B) and hippocampus (D) across age groups, with corresponding quantification of volume fraction. **(E, F)** Representative reconstructions of IBA1+ microglia in the cortex (E) and hippocampus (F) across the same age groups, with corresponding quantification of Sholl intersections, territory size, and soma-to-whole territory ratio. Age-group comparisons were performed relative to 4-month-old mice. Data are shown as mean ± SEM. Each symbol represents one analyzed cell. Scale bar represents 10 μm.

Microglia also demonstrated age-related morphological remodeling, although the timing varied by region. In the cortex, decline in Sholl intersections and territory size were apparent by 12 months and became more pronounced with advancing age (Fig. 3E). In the hippocampus, microglial morphology remained comparatively stable during midlife but again declined at 18 and 24 months (Fig. 3F). Both regions showed an age-associated increase in the soma-to-territory ratio, consistent with a shift toward a more compact, reactive morphology. In summary, these findings demonstrate that aging drives region-dependent remodeling of glial architecture. This late-life reduction in glial territory and process complexity offers a structural context for the age-associated decline in Cx43 observed in both the cortex and hippocampus.

### 2.4 Aging selectively enhances hemichannel activity while preserving gap junction communication in the hippocampus

To determine whether the observed Cx43 and neuroglial structural remodeling were accompanied by alterations in connexin channel function, we assessed hemichannel activity using ethidium bromide (EtBr) uptake in acute brain slices. In GFAP+ astrocytes, cortical EtBr uptake remained low across all age groups; in contrast, in hippocampus uptake increased significantly at 24 months; EtBr uptake is blocked by carbenoxolone (CBX), a broad Cx43 channel blocker (Fig. 4A). These results suggest that hemichannel activity is selectively upregulated in aged hippocampal astrocytes.

**Figure 4.**
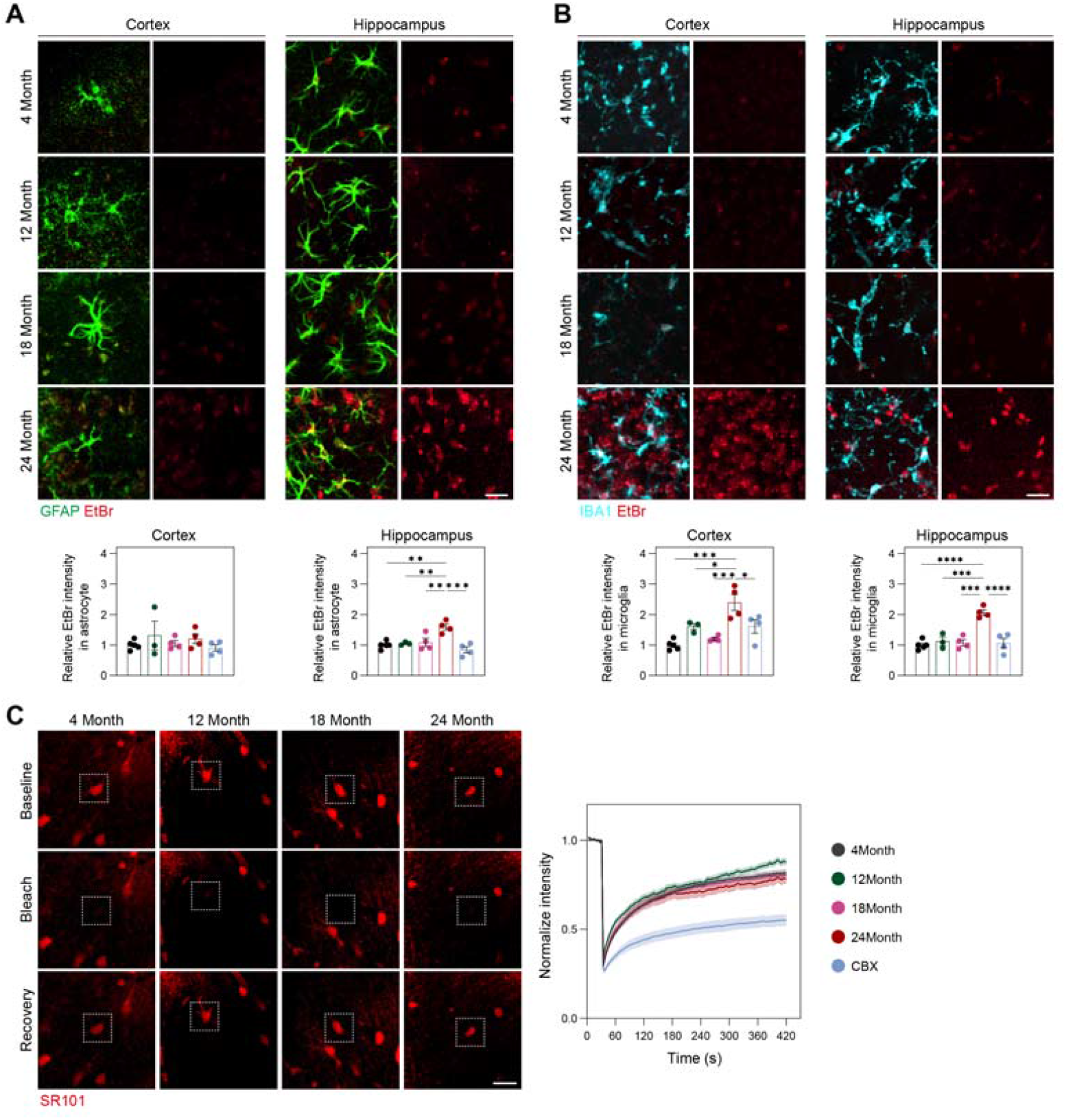
Connexin channel function is remodeled during brain aging. **(A)** Representative images and quantification of EtBr uptake in GFAP+ astrocytes from the cortex and hippocampus of mice aged 4, 12, 18, and 24 months. **(B)** Representative images and quantification of EtBr uptake in IBA1+ microglia from the cortex and hippocampus across the indicated ages. **(C)** Representative FRAP images of SR101-labeled tissue showing baseline, bleach, and recovery phases, together with normalized fluorescence recovery curves. Data are shown as mean ± SEM. Each dot represents one biological replicate. Scale bar represents 100 μm.

IBA1+ microglia displayed a more prominent advanced age-related increase in EtBr uptake. In both cortex and hippocampus, EtBr uptake remained relatively low until 18 months but increased markedly by 24 months. CBX treatment significantly inhibited EtBr uptake in microglia (Fig. 4B). Thus, while age-associated increases in microglial hemichannel activity occur in both regions, whereas such changes in astrocytes appear more region-specific.

We subsequently evaluated gap junction coupling in hippocampal astrocytes using fluorescence recovery after photobleaching (FRAP) assay, as previously described [20]. CBX treatment strongly suppressed fluorescence recovery, confirming the assay’s specificity for gap junction function. FRAP curves did not differ significantly between hippocampal astrocytes from 4-, 12-, 18-, and 24-month-old mice (Fig. 4C). This indicates that hippocampal astrocytic gap junction communication is preserved throughout the aging process.

In summary, aging remodels connexin-associated channel function in a highly selective manner. Rather than driving a broad decline in gap junction coupling, aging preferentially enhances hemichannel activity, particularly in microglia and hippocampal astrocytes at advanced age.

### 2.5 Human astrocytes show age-associated Cx43 reduction and structural remodeling

Given our findings in mice, we sought to determine whether astrocytic Cx43 remodeling is conserved in the aging human brain. To establish an independent, cell-type-resolved context, we interrogated the ZEBRA integrated brain atlas for *GJA1* expression in the human cortex and hippocampus [37]. Across all aging cohorts, *GJA1* was consistently detected in astrocytes, with lower or variable expression observed in microglia and vascular-associated cells (Fig. 5A). This validated astrocytes as the primary cell type for evaluating Cx43 remodeling in human cortical and hippocampal tissues.

**Figure 5.**
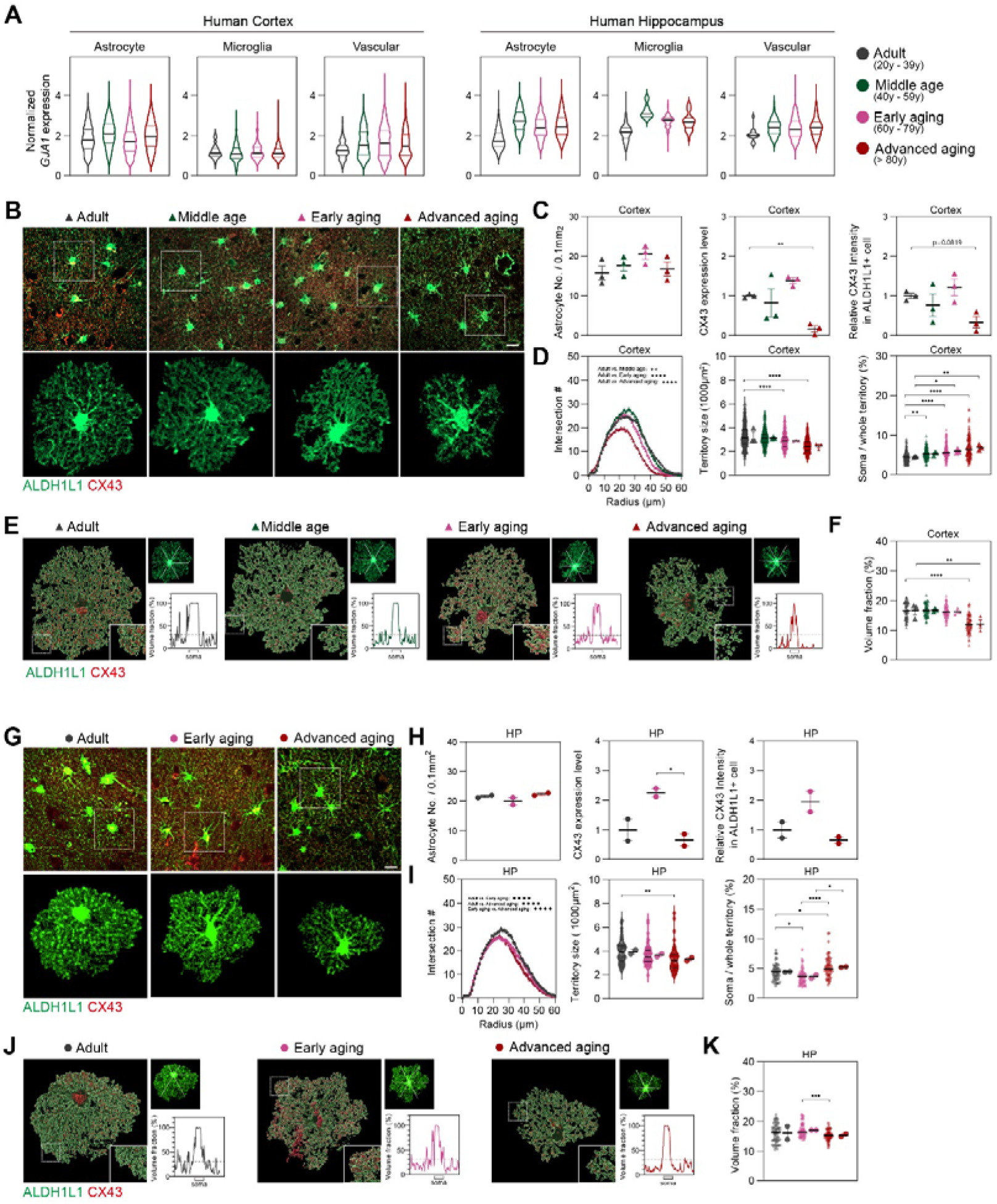
Astrocytic Cx43 expression and morphology are progressively remodeled in the aging human brain. **(A)** Normalized *GJA1* expression in astrocytes, microglia, and vascular cells from human cortex and hippocampus across adult, middle-age, early aging, and advanced-aging groups. **(B)** Representative images of ALDH1L1 and Cx43 immunostaining in human cortical tissue, together with representative reconstructions of individual astrocytes. **(C)** Quantification of astrocyte number, Cx43 expression level, and relative Cx43 intensity in ALDH1L1+ cells. **(D)** Sholl profiles and quantification of astrocyte territory size and soma-to-whole territory ratio. **(E)** Representative reconstructions illustrating subcellular Cx43 distribution within individual astrocytes across age groups. **(F)** Quantification of Cx43 volume fraction within reconstructed astrocytes. Data are shown as mean ± SEM. Each symbol represents one biological sample or analyzed cell. Scale bar represents 20 μm.

In the human cortex, the density of ALDH1L1+ astrocytes remained stable across adult, middle-aged, early aging, and advanced-aging groups (Fig. 5B). However, Cx43 expression exhibited a progressive decline in advanced aging, characterized by reduced tissue-level signal and a concurrent decrease in intensity within ALDH1L1+ astrocytes (Fig. 5C). This molecular remodeling coincided with pronounced structural simplification; 3D reconstruction and Sholl analysis revealed reduced branching complexity, contracted territorial domains, and an increased soma-to-whole territory ratio reflecting age-dependent astrocyte atrophy (Fig. 5D). Furthermore, volume fraction analysis confirmed a reduction in fine-process-rich domains, indicating that aging disrupts both major branches and the fine-process architecture of human astrocytes (Fig. 5E, F).

A comparable pattern was evident in the human hippocampus. While astrocyte density remained unchanged, Cx43 expression decreased significantly in advanced aging (Fig. 5G, H). Hippocampal astrocytes also exhibited atrophy as indicated by reduced process complexity, territorial contraction, and an increased soma-to-whole territory ratio (Fig. 5I), paralleling the morphological shifts observed in the cortex. Additionally, volume fraction analysis confirmed a decline in fine-process-rich astrocytic domains with age (Fig. 5J, K).

Collectively, these data demonstrate that human astrocytes undergo coordinated Cx43 and morphological remodeling during aging. Consistent decline in Cx43 expression, coupled with branch simplification and the loss of fine processes, mirrors our observations in mice. This suggests that aging promotes the structural and molecular reorganization of the astrocytic connexin-mediated network, rather than simply inducing a loss of astrocyte abundance.

### 2.6 Human transcriptomic analyses link *GJA1* remodeling to region-dependent astrocyte state organization

To investigate whether Cx43/*GJA1* remodeling correlates with astrocyte functional states in the aging human brain, we analyzed human prefrontal cortex (PFC) and hippocampal single-cell datasets [38, 39]. We calculated six astrocyte modules representing homeostatic support, metabolic support, BBB interaction, cell morphology, reactive state, and inflammatory-related programs (Table 1); notably, *GJA1* was excluded from the gene sets defining these modules.

**Table 1.**
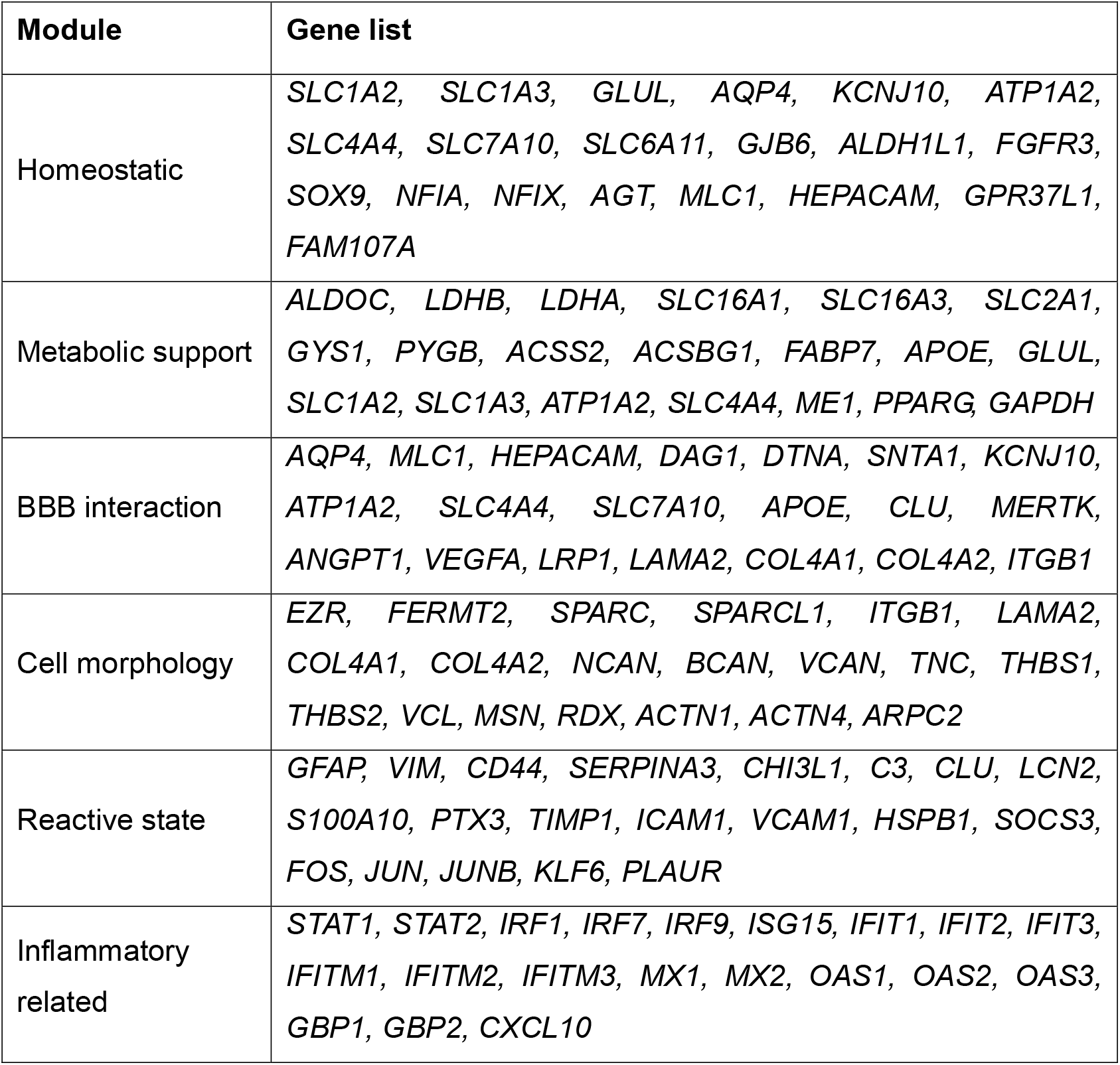
Genes used for module score analysis.

At the sample level, chronological age, when analyzed independently, showed no strong monotonic association with most astrocyte module scores in either the PFC or hippocampus (Fig. 6A, C). These results suggest that aging astrocytes largely retain their core transcriptional program structure, rather than undergoing a broad transcriptomic decline. The observed age-related changes were modest and region-dependent, for example, astrocytes in the PFC exhibited a modest negative trend in the metabolic-support module, whereas hippocampal astrocytes showed weak negative trends in the BBB interaction and cell morphology modules. These trends suggest that there is no uniform, age-related linear shift in astrocyte functional module activity.

**Figure 6.**
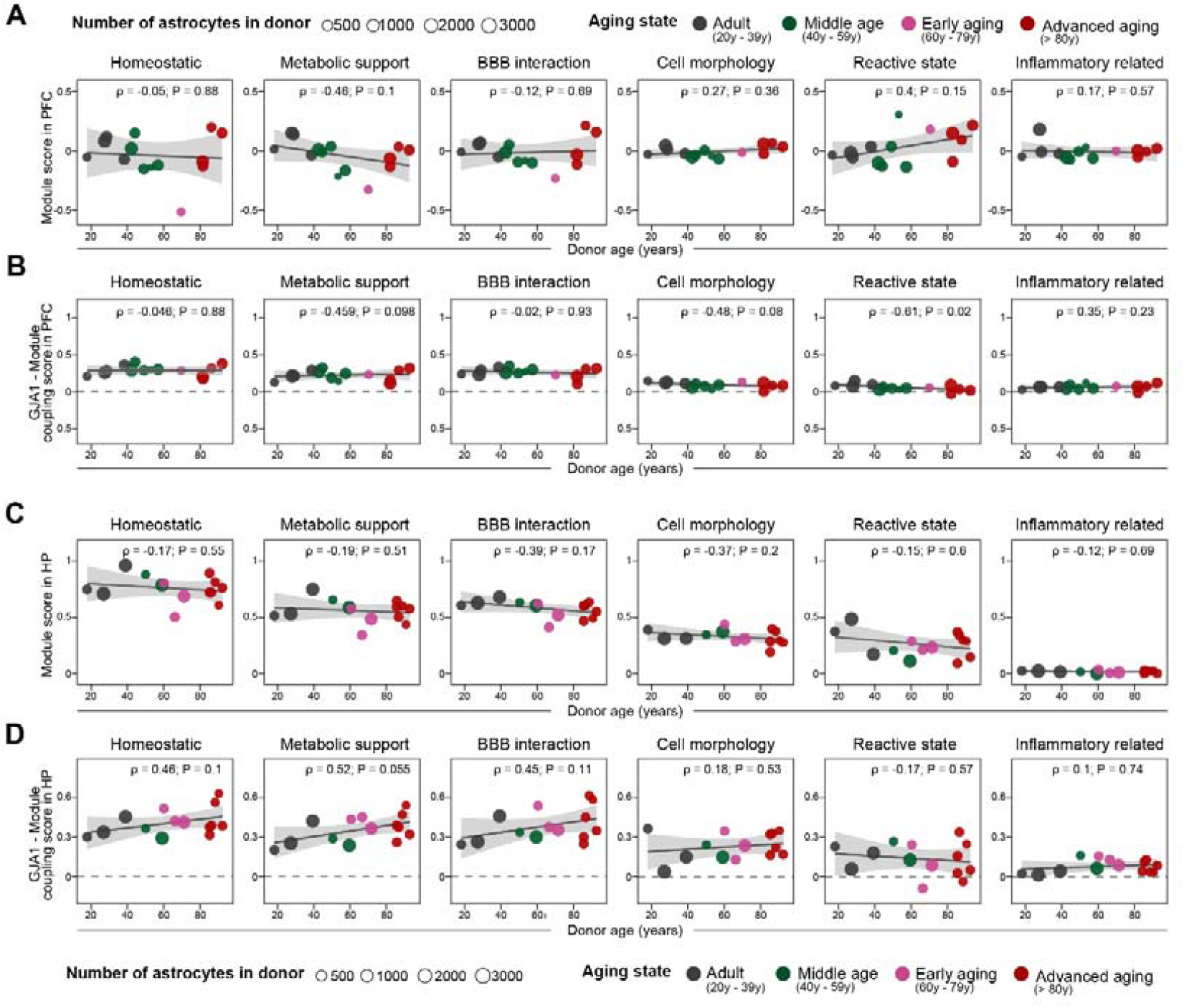
Human PFC and hippocampal transcriptomes reveal region-dependent *GJA1*-associated astrocyte state organization during aging. Donor-level analysis of astrocyte functional module scores across age in human prefrontal cortex astrocytes. **(B)** Age-related trends in *GJA1*-module coupling in PFC astrocytes. For each donor, *GJA1*-module coupling was calculated as the within-donor Spearman correlation between *GJA1* expression and each 20-gene module score. **(C)** Donor-level analysis of astrocyte functional module scores across age in human hippocampal astrocytes. **(D)** Age-related trends in *GJA1*-module coupling in hippocampal astrocytes. Within-donor *GJA1*-module coupling was calculated as in panel B. Each point represents one donor. Point size indicates the number of astrocytes included for that donor. Gray lines indicate fitted trends, and shaded bands indicate 95% confidence intervals. Spearman correlation coefficients and P values are shown in each panel.

We next evaluated whether the coupling between *GJA1* expression and these modules changed with age. Age alone did not strongly increase any single astrocyte module. Instead, aging changed the coupling between *GJA1* and selected modules in a region-dependent manner. In PFC astrocytes, the link between *GJA1* and the reactive-state module declined with age, with a similar negative trend observed for the cell structural module (Fig. 6B). Conversely, in hippocampal astrocytes, *GJA1* coupling with the homeostatic, metabolic-support, and BBB interaction modules showed positive age-related trends (Fig. 6D). These results indicate that *GJA1*-associated astrocyte transcriptional states undergo region-dependent reorganization during aging.

In conjunction with our human histological findings, these data support the interpretation that Cx43/*GJA1* remodeling corresponds to the reorganization of astrocyte functional states in the aging human brain. This remodeling does not appear to reflect a global loss of astrocytes or a uniform alteration in all astrocytic functional programs. Instead, our findings point to region-specific changes in the link between *GJA1* and astrocyte programs related to cellular support, morphology, and reactive transcriptional states.

## 3. DISCUSSION

This study characterizes the multifaceted remodeling of astrocytic Cx43 during aging. Our results reveal a non-linear, region-dependent trajectory of Cx43 expression rather than a uniform age-related decline in both humans and mice. This molecular remodeling coincides with glial structural simplification and selective alterations in Cx43 hemichannel activity, demonstrating that brain aging drives a comprehensive reorganization of astrocytic connexin biology across molecular, structural, and functional levels.

A central implication of our findings is that Cx43 regulation during aging constitutes dynamic remodeling rather than simple protein loss. Cx43 mediates diverse functions, including gap junction coupling, hemichannel activity, and non-channel signaling [24, 40-43]. The observed non-linear expression pattern argues against a sole age-related reduction, instead suggesting a stage-dependent trajectory that may reflect adaptive or compensatory responses.

The region-dependent pattern of Cx43 remodeling aligns with the broader understanding of brain aging. Recent single-cell and spatial studies have demonstrated that aging involves highly specific transcriptional changes across brain regions and cell types, including modifications in glial cells, vascular-associated cells, and neuronal support programs [38, 44].Our data extend this concept to astrocytic Cx43. In both the mouse cortex and hippocampus, Cx43 signal intensity varied with age while astrocyte number remained stable. The most evident decline in Cx43 signal and astrocyte structural complexity occurred at advanced age. These findings indicating that the aging phenotype is not driven by astrocyte loss, but rather by remodeling within glial-neuronal network.

Astrocyte structural remodeling provides important context for interpreting these Cx43 changes. Astrocytes support neuronal and vascular function within their territories formed by major branches, fine processes, and endfeet associated with the vasculature [9, 12]. In our study, aging reduced complexity of astrocyte branching, territorial coverage, and fine processes in both mouse and human brain. These changes may reduce astrocytic interaction with synapses, vessels, and neighboring glial cells. Therefore, the late-life reduction in astrocyte Cx43 may reflect a broader remodeling of astrocyte architecture and local homeostatic capacity, rather than an isolated change in connexin abundance. Microglia also showed age-associated structural remodeling. In both cortex and hippocampus, aging was associated with reduced process complexity and a shift toward a more compact morphology, which is consistent with the known age-dependent changes in microglia [38, 39]. Our data show that microglial morphology did not fully mirror astrocyte morphology in timing or regional pattern. This supports the idea that astrocytes and microglia follow related yet distinct aging trajectories, and suggests that Cx43-associated aging should be interpreted within a broader glial remodeling framework, rather than only as an astrocyte-intrinsic event.

Our functional data demonstrate that Cx43 abundance and connexin channel activity are distinct parameters during aging. While astrocytic Cx43 levels declined in advanced age, hippocampal astrocytic gap junction, as measured by FRAP, remained preserved. In contrast, EtBr uptake increased in aged hippocampal astrocytes and microglia across the cortex and hippocampus. These results suggest that aging preferentially enhances hemichannel activity in specific glial populations, rather than precipitating a broad collapse of astrocytic gap junction coupling. This distinction is critical, as gap junction channels and hemichannels mediate distinct functions. While gap junction facilitates direct intercellular exchange to maintain the brain homeostasis, whereas hemichannels support molecular exchange between the cytoplasm and the extracellular space [24]. Under inflammatory or stress conditions, sustained hemichannel opening can become harmful by promoting ATP, glutamate, and danger-signal release, thereby amplifying extracellular inflammatory signaling and local tissue stress [27]. Therefore, the age-associated increase in hemichannel dye uptake may represent a potentially component of glial aging, even when gap junction coupling is preserved.

Our human data corroborate the relevance of this remodeling process beyond the mouse model. In both the human cortex and hippocampus, astrocyte density maintained, whereas astrocyte Cx43 declined during advanced aging. Human astrocytes exhibited reduced branching complexity, contracted territorial domains, higher soma-to-territory ratios, and a loss of fine-process-rich domains. These findings align with our mouse data, suggesting that astrocytic Cx43 remodeling is evolutionarily conserved. This interpretation is consistent with previous human aging studies demonstrating that astrocytes undergo significant molecular and structural reorganization even in the absence of substantial cell loss [18, 38].

Furthermore, our human single-cell analyses provided a transcriptomic dimension to this model. Age alone did not drive strong monotonic shifts in curated astrocyte functional modules in either the PFC or hippocampus, suggesting that global astrocyte module activity is not strictly determined by chronological age. Nevertheless, the relationship between *GJA1* expression and astrocyte functional programs exhibited distinct, region-dependent, age-related patterns. In PFC astrocytes, *GJA1* coupling with reactive-state and cell-structural modules tended to weaken with age. Conversely, in hippocampal astrocytes, *GJA1* coupling with homeostatic, metabolic-support, and BBB-interaction modules exhibited positive age-related trends. These findings suggest that *GJA1*-associated astrocyte states undergo reorganization during aging in a region-specific manner. This observation is consistent with established regional heterogeneity in astrocytes and supports the hypothesis that cortical and hippocampal astrocytes follow distinct aging trajectories [17, 45]. Collectively, these transcriptomic findings aid in interpreting our tissue-level results. The reduction in Cx43 signal and astrocyte structural complexity in advanced age does not necessarily imply a global loss of astrocyte function; rather, it may reflect a reorganization of astrocyte states. Within this framework, *GJA1* should not be considered a singular causal driver of aging, but rather an astrocyte-associated molecular axis linked to broader functional adaptive programs.

While our cross-sectional analyses offer a robust framework, future longitudinal studies are required to disentangle whether these changes reflect adaptive compensation during physiological aging. In summary, our study identifies Cx43 remodeling as a distinct molecular, structural, and functional hallmark of brain aging. Across both mouse and human models, aging was consistently associated with altered astrocytic Cx43 level, glial structural simplification, and selective modifications in Cx43 channel function. Furthermore, our human single-cell analyses indicate that *GJA1*-associated transcriptional programs are reorganized in a region-specific manner. Collectively, these findings support a model wherein aging reshapes astrocytic connexin biology and local glial homeostasis, potentially establishing a brain environment more vulnerable to subsequent neurodegenerative stress.

## 4. METHODS

### Animals

Mice used in this study were obtained from Zhuhai BesTest Bio-Tech Co,.Ltd, and maintained in a pathogen-free animal facility with 12 h/12 h light/dark cycle under suitable temperature and humidity. All animal experimental procedures were conducted in accordance with the ethical approval of the Sun Yat-Sen University Institutional Animal Care and Use Committee. For sample preparation, mice were anesthetized by intraperitoneal injection of 1.25% avertin at 20 μL/g, followed by ice cold PBS transcranial perfusion. The brain was then isolated, the left-half was dissection to different brain regions and snap-frozen at -80 °C, while the right-half was subjected to post-fixation with 4% PFA at 4 °C for 24 h and following immersed in 30% sucrose prepared for cryoprotection.

### Human sample

Human post-mortem brain tissues were provided by the Human Brain Bank, Chinese Academy of Medical Sciences & Peking Union Medical College (Beijing, China) under the ethical approval by the Institutional Review Board of the Institute of Basic Medical Sciences, Chinese Academy of Medical Sciences (Approval Numbers: 009-2014 and 2022125). Informed consent was obtained by participants.

### Immunostaining

For mice, brain slices were cryostat sectioning to 20 μm thickness. Sections were processed for immunostaining by overnight incubation at 4□°C with primary antibodies, followed by Alexa Fluor conjugated secondary IgG antibodies for 2 h at room temperature. Then, the sections were washed, mounted, and examined with slide scanner or confocal microscope. For the quantification of sections, five brain slices from each mouse brain were analyzed. Fiji software was used to quantify the fluorescent intensity, which was averaged for each mouse and then normalized to control group as relative change.

For human brain sample, after deparaffinization, the paraffin-embedded sections were treated with 3% H_2_O_2_ and processed for antigen retrieval. The sections were blocked with blocking solution and incubated with targeted primary antibody for 2 h at 37 °C and overnight at 4 °C, then followed by Alexa Fluor conjugated secondary IgG antibodies for 2 h at room temperature. The stained sections were examined with confocal microscope. For the quantification, we randomly selected six non-overlapping areas of 1 mm^2^ in the cortex and hippocampus, respectively. Fiji software was used to quantify the fluorescent intensity in each image; then these intensities were averaged for each human brain. Finally, the fluorescent intensities were normalized to the controls.

### Sholl analysis

Sholl analysis was performed to quantify the morphological complexity of individual astrocytes and microglia as previously described [27]. Confocal images of ALDH1L1+ astrocytes and IBA1+ microglia were acquired from the cortex and hippocampus of mice at different ages. Individual cells with clearly identifiable soma and processes, minimal overlap with neighboring cells, and complete morphology within the imaging field were selected for analysis. Images were converted to 8-bit grayscale images and processed in ImageJ. For each cell, the soma was manually defined as the center point, and concentric circles were generated at fixed radial intervals from the soma. The number of intersections between cellular processes and each concentric circle was quantified to generate the Sholl profile. The maximum number of intersections and the distribution of intersections across radial distances were used to assess process complexity. Territory size was measured as the projected area occupied by the reconstructed cell, and the soma-to-whole territory ratio was calculated by dividing the soma area by the total cellular territory area.

### Volume fraction analysis

Volume fraction analysis was performed to evaluate the proportion of process-rich domains within the territory of individual astrocytes, as previously described [19]. Confocal images of ALDH1L1+ astrocytes in the cortex and hippocampus of mice at different ages were acquired under identical experimental conditions and using consistent imaging parameters. Individual astrocytes with clearly identifiable soma and well-defined territorial boundaries were selected for analysis. After background subtraction, images were converted to 8-bit grayscale images in ImageJ. The astrocytic territory was manually delineated according to the outer boundary of ALDH1L1+ processes. Using the straight-line tool in ImageJ, three lines were drawn through the soma at 60° intervals, with the soma center defined as the origin. Fluorescence intensity profiles along each line were extracted using the Plot Profile function, generating six semi-profiles for each cell. For each profile, the peak fluorescence intensity within the soma was used as the reference value and defined as 100% volume fraction (VF). VF was calculated as follows: VF (%) = (F_local / F_soma) × 100, where F_local represents the local fluorescence intensity along the line profile and F_soma represents the peak fluorescence intensity within the soma. The fluorescence intensity within the astrocytic territory was normalized to the somatic reference value to estimate the relative volume fraction of ALDH1L1+ astrocytic structures. For subcellular distribution analysis, linear fluorescence intensity profiles extending from the soma to the distal territory were plotted as normalized fluorescence intensity against distance from the soma. The mean volume fraction within each astrocytic territory was used for statistical comparison among age groups.

### Acute brain slices preparation

Acute brain slices were prepared following the previously published protocols [46]. Fresh brains from mice of different age were rapidly removed and immersed in chilled NMDG-HEPES artificial cerebrospinal fluid (aCSF) for 1 min before slicing. Slices were then transferred to NMDG-HEPES aCSF maintained at 34 °C. Na^+^ spike-in solution was added every 2 min for a total of 10 min. The slices were subsequently incubated in HEPES aCSF for 1 h and then maintained in oxygenated recording aCSF before experiments. All aCSF solutions used for slice recovery and recording were continuously bubbled with 95% O_2_ and 5% CO_2_ unless otherwise stated.

### Gap Junction Fluorescence Recovery After Photobleaching (gap-FRAP) assay

Gap junction coupling was assessed using a fluorescence recovery after photobleaching (FRAP) assay as previously described [47]. Acute brain slices were incubated with 50 µM soulforodamin 101 (SR101) for 30 min at 34 °C and then transferred to the recording chamber of an FV3000 confocal microscope (Olympus). Baseline fluorescence was acquired at low laser power every 5 s for 10 cycles over a 158 × 158 µm field of view. A single target astrocyte was then photobleached using high laser power over a 5 × 5 µm region for 20 s. Fluorescence recovery was recorded for 7 min after photobleaching. Fluorescence intensity at each time point was normalized to F_0_, defined as the mean baseline fluorescence intensity before bleaching, and expressed as percentage recovery relative to baseline fluorescence intensity (F□/F_0_ × 100%).

### Dye uptake

Dye uptake in acute brain slices was measured according to previously published protocol, with minor modifications [26]. Briefly, acute brain slices were preincubated in calcium-free aCSF, standard aCSF, or aCSF containing the hemichannel blocker carbenoxolone (CBX, 200 µM) for 15 min at room temperature. Brain slices were then incubated with ethidium bromide (EtBr, 4 µM) for 10 min at room temperature. After dye loading, slices were fixed in 4% paraformaldehyde for 1 h and then processed for immunofluorescence staining.

For immunofluorescence staining, slices were permeabilized with 1% Triton X-100 in PBS for 30 min and blocked with 2% BSA. Slices were incubated with primary antibodies overnight at 4 °C, followed by incubation with the corresponding secondary antibodies. Images from the target brain regions were acquired using an IXplore SpinSR confocal microscope (Olympus). To ensure consistency across samples, images were collected from the same depth below the slice surface, approximately 10 µm, using identical imaging intervals and acquisition settings.

### Bioinformatic analysis of public datasets

Publicly available aging-related transcriptomic, proteomic, and single-cell datasets were re-analyzed to examine the tissue distribution, cellular enrichment, and age-associated transcriptional organization of Cx43/*GJA1*. For cross-tissue analyses, published multi-organ aging proteomic and transcriptomic datasets were used to compare Cx43 protein abundance and *Gja1* mRNA expression across organs and ages. Expression values were extracted from the processed matrices provided by the original studies and visualized across organs or age groups without additional batch correction.

For cell-type enrichment analyses, published mouse and human brain single-cell or single-nucleus RNA-seq atlases were used Cells were grouped according to the major cell-type labels provided by the original studies, including astrocytes, microglia, oligodendrocyte-lineage cells, vascular-associated cells, and neuronal populations. *Gja1* or *GJA1* expression was then summarized across annotated cell types to determine the major cellular compartments associated with Cx43 expression in the brain.

For human aging analyses, prefrontal cortex and hippocampal single-cell or single-nucleus transcriptomic datasets were analyzed separately. Cells annotated as astrocytes were retained for downstream analysis. Sample metadata, including age and sample identity, were extracted from the original metadata files. Samples were grouped into age stages when required for visualization. The age-stage categories were defined as adult, middle age, early aging, and advanced. Astrocyte transcriptional programs were evaluated using curated gene modules representing homeostatic support, metabolic support, vascular/BBB interaction, structural or morphology-related features, reactive/stress-response programs, and inflammatory-related programs. These modules were used as continuous transcriptional program scores rather than mutually exclusive astrocyte-state labels. *GJA1* was excluded from all module gene sets to avoid circular correlation between *GJA1* expression and module scores. Module scores were calculated at the single-cell level using normalized expression data. For each sample, astrocyte module scores were then averaged to generate sample-level module scores. Sample-level associations between chronological age and module activity were assessed using Spearman correlation analysis. To examine the relationship between *GJA1* and astrocyte transcriptional programs, within-sample correlations between *GJA1* expression and each module score were calculated across astrocytes from the same sample. These correlation coefficients were then used as sample-level measures of *GJA1*–program coupling.

All bioinformatic analyses were performed in R. Single-cell data processing, subsetting, normalization, and module scoring were performed using Seurat. Data manipulation and visualization were performed using standard R packages.

### Statistical analysis

Statistical significance between groups was determined with GraphPad Prism software. The unpaired t test was used to determine the difference between the two groups. One-way analysis of variance (ANOVA) was used to determine the difference among three and four groups. Two-way ANOVA was used to determine the difference between group variables in the curves. The Pearson correlation test was used to determine the correlation between variables. Data distribution was assumed to be normal, but this was not formally tested. A probability of p < 0.05 was considered statistically significant. All significant statistical results are indicated within the figures with the following conventions: *p < 0.05, **p < 0.01, ***p < 0.001, ****p < 0.0001. Error bars represent mean ± standard deviation. No statistical methods were used to predetermine sample sizes. The sample size per group was determined from previous publications using a similar methodology. Investigators were blinded to group allocation during data analysis. All experiments were performed at least three times, and the findings were replicated in individual mice in each experiment.

## ACKNOWLEDGMENTS

This work was supported by National Natural Science Foundation of China (32400809 to Q.W.), Shenzhen Fundamental Research Program (RCJC20231211090018040 and ZDSYS20220606100801003 to C.Y.).

Human tissue was provided by the National Human Brain Bank for Development and Function, Chinese Academy of Medical Sciences and Peking Union Medical College, Beijing, China. This study was supported by the Institute of Basic Medical Sciences, Chinese Academy of Medical Sciences, Neuroscience Center, and the China Human Brain Banking Consortium, by grants from STI2030-Major Project #2021ZD0201100 Task 1 #2021ZD0201101. We thank Bioimaging Core of Shenzhen Bay Laboratory for providing imaging support. We also would like to acknowledge Bioimaging Core engineer Lan Yuan for assistance with the SpinSR spinning disk confocal microscopy.

## Author contributions

Q.W., C.W., and H.L. conducted most of the experiments and wrote the original draft. M.S. and W.J. contributed to human samples collection. Y.C. and A.V. conceptualized the study, contributed to the interpretation of the results, and edited and reviewed the manuscript.

## DATA AVAILABILITY STATEMENT

All data that support the findings of this study are available from the corresponding author upon reasonable request.

